# Disintegrin-like and Cysteine-rich Domains Govern Enzymatic Activity and Substrate Recognition in *Echis* Snake Venom Metalloproteinases

**DOI:** 10.64898/2026.08.26.747306

**Authors:** Sophie Hall, Bronwyn Rand, Iara Aimê Cardoso, Adam Robinson, Mark C. Wilkinson, Dakang Shen, Sebastian Fernandez, Georgia Balchin, Konrad Kamil Hus, Alastair W. Poole, Nicholas R. Casewell, Imre Berger, Christiane Schaffitzel

## Abstract

Snake venom metalloproteinases (SVMPs) are major drivers of pathology following viper envenomation and represent important targets for the development of next-generation recombinant antivenoms. PIII SVMPs are among the most potent haemorrhagic toxins and contain disintegrin-like (Dis) and cysteine-rich (C-rich) accessory domains. Despite their biomedical importance, the mechanistic roles of these accessory domains in substrate recognition and catalysis remain poorly understood. We produced recombinant full-length and domain-deletion variants of two functionally distinct PIII SVMPs: the broadly proteolytic, cytotoxic cPIII and the highly specific prothrombin activator Ecarin. Proteins were expressed as latent zymogens in insect cells, auto-activated by Zn²⁺, and analysed using enzymatic, blood clotting, and cell-based assays. Progressive removal of the C-rich and Dis domains reduced zymogen auto-activation and markedly diminished catalytic activity in both toxins. In cPIII, domain deletion caused a stepwise loss of proteolytic and cytotoxic activity without altering substrate preference. In Ecarin, removal of the accessory domains strongly impaired prothrombin activation, and thus plasma clotting, demonstrating a critical role in substrate recognition. Conversely, deletion of the C-rich domain increased fibrinogenolytic activity, revealing a substrate-dependent gatekeeping function. Deglycosylation showed that N-linked glycans modulate SVMP activity in a construct-dependent manner. Recombinant Ecarin closely recapitulated the biochemical properties of the native venom-derived toxin. Our data support a model in which PIII SVMP accessory domains enhance substrate positioning and catalytic efficiency while selectively restricting access to non-cognate substrates. These findings establish accessory-domain-mediated substrate recognition as a key determinant of SVMP function, informing rational antivenom design.

## INTRODUCTION

Snake venom is a complex cocktail of more than 100 proteins and peptides (Oliveira *et al*, 2022). Among the most abundant and medically significant components are snake venom metalloproteinases (SVMPs), zinc-dependent enzymes that contribute to haemorrhage, coagulopathy, and tissue necrosis following envenomation (Bittenbinder *et al*, 2024). SVMPs belong to the metzincin superfamily (Bastos *et al*, 2016) and are especially expanded and diversified in viperid venoms, where they represent major drivers of venom-induced pathology (Dawson *et al*, 2021; Fox & Serrano, 2008; Gutierrez & Rucavado, 2000; Olaoba *et al*, 2020).

SVMPs have evolved via gene duplication events followed by extensive domain loss, resulting in a structurally and functionally diverse modular enzyme family (Casewell *et al*, 2011). They are classified into three major groups with different domain composition: PI SVMPs, which consist solely of the metalloproteinase (MP) domain, PII SVMPs, which additionally contain a disintegrin domain involved in integrin binding and modulation of cell-matrix interactions, and PIII SVMPs, which comprise a metalloproteinase domain followed by a disintegrin-like (Dis) and a cysteine-rich (C-rich) domain (Olaoba *et al*., 2020). This modular architecture contributes to substantial variation in substrate specificity, tissue targeting, and toxic potency (Olaoba *et al*., 2020; Takeda *et al*, 2012).

SVMP domains are stabilised by extensive disulphide bonding, with proteins such as Ecarin, a PIII SVMP, containing 38 cysteine residues (Nishida *et al*, 1995). In PIII SVMPs, the C-rich domain is positioned adjacent to the catalytic cleft of the MP domain and has been implicated in substrate recognition and binding, although its precise mechanistic role remains elusive. Two non-mutually exclusive functional models were proposed to explain the function of the C-rich domains in SVMPs and in the related ADAM (A Disintegrin And Metalloproteinase) family (Gaultier *et al*, 2002; Serrano *et al*, 2006): a selective ‘recruitment-feeding’ mechanism in which accessory domains cooperate to capture and guide substrates into the catalytic cleft and a ‘gatekeeping’ mechanism in which the domain restricts access of non-cognate substrates to the active site (Figure 1a).

**Figure 1.**
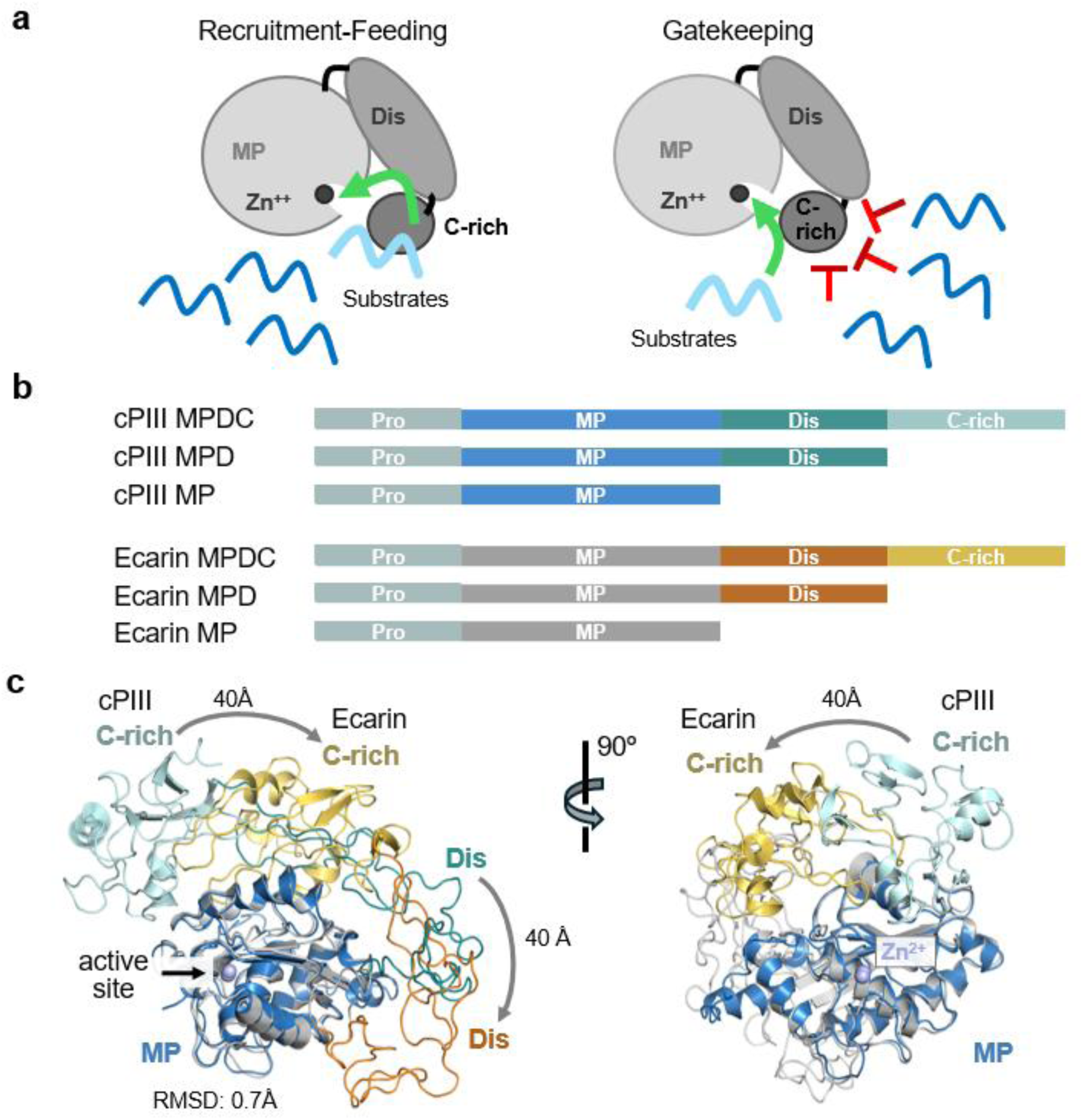
SVMPs are multidomain metalloproteinases. (a) Two proposed mechanisms for the C-rich domain in regulating access to the active site. Target substrates are shown in light blue, non-target proteins in dark blue. SVMPs are show schematically. Metalloproteinase (MP), Dis and C-rich domains are indicated. (b) Simplified schematic of the constructs used in this study: cPIII: a cytotoxic PIII SVMP fro*m Echis carinatus sochureki* (UniProt E9JG34), and Ecarin from *Echis pyramidum leakeyi* (UniProt Q90495). (c) Alignment of the MP domains (RMSD 0.7 Å) of AlphaFold 3 predicted model of cPIII (MP, Dis and C-rich domains shown in blue, teal and cyan respectively) and the Ecarin structure (PDB: 9CLP) (MP, Dis, C-rich domains shown in grey, orange and yellow respectively). Differences in the positioning (40Å) of the C-rich and desintegrin domain in Ecarin as compared to cPIII is indicated by curved arrows. Structural alignment and visualisation were performed in PyMOL.

To dissect the role of the Dis and C-rich domains, we used our baculovirus/insect cell expression system, which enables the production of correctly folded, post-translationally processed, recombinant SVMPs at milligram scale (Bieniossek *et al*, 2012; Hall *et al*, 2026). This recombinant approach provides a significant advantage over purification from native venom, where isolating specific isoforms is often limited by their low abundance, heterogeneity, and loss of activity during purification (Hall *et al*., 2026). Our recombinant approach further allows the functional comparison of PI-like, PII-like and PIII SVMPs with the same metalloproteinase domain, which is not possible with native venom-derived toxins.

SVMPs were expressed as latent zymogens using their native prodomain, which directs secretion and maintains the enzyme in an inactive state. Latency is enforced by a conserved propeptide, which is part of the prodomain and occupies the catalytic cleft (Fox & Serrano, 2005; Moura-da-Silva *et al*, 2016). The propeptide coordinates the active-site Zn²⁺ ion via a cysteine residue, consistent with a cysteine-switch-like inhibitory mechanism (Grams *et al*, 1993). This inhibition is stabilised by the prodomain scaffold, which positions the propeptide across the active site and prevents substrate access (Grams *et al*., 1993). SVMP activation occurs by proteolytic removal of the prodomain, exposing the catalytic site with a Zn^2+^ ion coordinated by three conserved histidine residues which are essential for catalysis (Grams *et al*., 1993; Hall *et al*., 2026). Our strategy enables production of cytotoxic SVMPs as zymogens, thus not killing the cells that produce them, and their auto-activation into a mature form after purification, by incubating with Zn^2+^ ions.

Here, we studied the impact of the accessory Dis and C-rich domains on the proteinase activity of two functionally distinct PIII SVMPs by generating deletion constructs (Figure 1b). First, we analysed a highly toxic PIII SVMP (cPIII) from *Echis carinatus sochureki* (UniProt: E9JG34), which exhibits broad substrate specificity, including fibrinogenolytic, caseinolytic, and insulin-degrading activity, as well as strong activity toward the fluorogenic substrate ES010 (Hall *et al*., 2026). Second, we expressed and purified the procoagulant PIII SVMP Ecarin from *Echis pyramidum leakeyi* (UniProt Q90495), which displays a very narrow substrate spectrum and specifically activates prothrombin to thrombin via the intermediate meizothrombin, leading to rapid blood clot formation (Francischetti & Reyes Gil, 2019). Structural comparison of these two PIII SVMPs indicates variability in the relative orientation of the Dis and C-rich domains compared to the active site (Figure 1c), which may contribute to the observed differences in substrate specificity and functional activity. Using the recombinant full-length and truncated SVMP constructs, we show that the Dis and C-rich domains of PIII SVMPs are key determinants of catalytic activity and substrate specificity. Removal of the Dis and C-rich domains strongly reduces proteinase activity and affects cleavage across fluorogenic, chromogenic, and physiological substrates, supporting a substrate feeding mechanism. Deletion of the C-rich domain of Ecarin also appears to affect substrate selection, leading to cleavage of fibrinogen which full-length Ecarin does not recognise, supporting a gatekeeping mechanism. Our systematic study provides strong evidence that Dis and C-rich domains enhance the catalytic activity of the MP domain and mediate substrate recognition.

## RESULTS

### Design of domain deletion constructs from functionally divergent PIII SVMPs

Our study is based on two well-characterised model PIII SVMPs with very different substrate spectra and toxicity. We chose the cytotoxic PIII (cPIII) from *Echis carinatus sochureki* (UniProt ID: E9JG34) (Hall *et al*., 2026) and haemotoxic Ecarin from Kenyan *Echis pyramidum leakeyi* (UniProt ID: Q90495) (Jafari *et al*, 2022) (historically classified as *Echis carinatus*), which specifically activates prothrombin leading to blood clotting (Misson Mindrebo *et al*, 2024). Sequence alignment of cPIII and Ecarin (Supplementary Figure 1) shows 65% identity between the two sequences, highlighting the high conservation of the prodomain and the active site of the metalloproteinase domain. In contrast, the sequences of the Dis and C-rich domains are strongly divergent. To investigate the contributions of the Dis and C-rich domains to metalloproteinase activity and substrate selection, we designed cPIII and Ecarin SVMPs with these domains removed (Figure 1b). The boundaries of MP, Dis and C-rich domains were taken from UniProt and verified using structural models predicted by AlphaFold3 (Figure 1c) (Abramson *et al*, 2024). The resulting constructs differ in domain complexity and the number of cysteine residues, which may influence protein folding, aggregation, activation efficiency and proteinase activity. The full-size cPIII MPDC zymogen contains four domains (prodomain, metalloproteinase domain (MP), Dis (D) and C-rich (C) domain) with 37 cysteines, cPIII MPD (PII-like) has three domains with 24 cysteines and cPIII (PI-like) MP has two domains with 9 cysteines. Similarly, Ecarin MPDC comprises four domains and 38 cysteines, MPD (PII-like) has three domains and 25 cysteines, and MP (PI-like) consists of two domains with 10 cysteines (Figure 1b).

### Production of full-length PIII SVMP and deletion constructs

Each SVMP expression construct was designed as a zymogen with a native signal peptide preceding the N-terminal prodomain. The prodomain maintains enzyme latency during expression, considerably reducing the cytotoxicity of the SVMPs (Hall *et al*., 2026). C-terminal His- and Avi-tags were added for identification, purification and other downstream processes (Hall *et al*., 2026). The DNA constructs were codon optimised for *Spodoptera frugiperda* (Sf21) and *Trichoplusia ni* (Hi5), and SVMPs were expressed as secreted proteins in Hi5 cells using the MultiBac baculovirus/insect cell expression system (Bieniossek *et al*., 2012).

Following secretion of SVMPs into the culture media, a three-step purification process was performed consisting of immobilised metal affinity chromatography (IMAC), anion exchange chromatography (IEX) and size exclusion chromatography (SEC), where they generally eluted in one prominent peak mostly as monomeric proteins (Figure 2).

**Figure 2:**
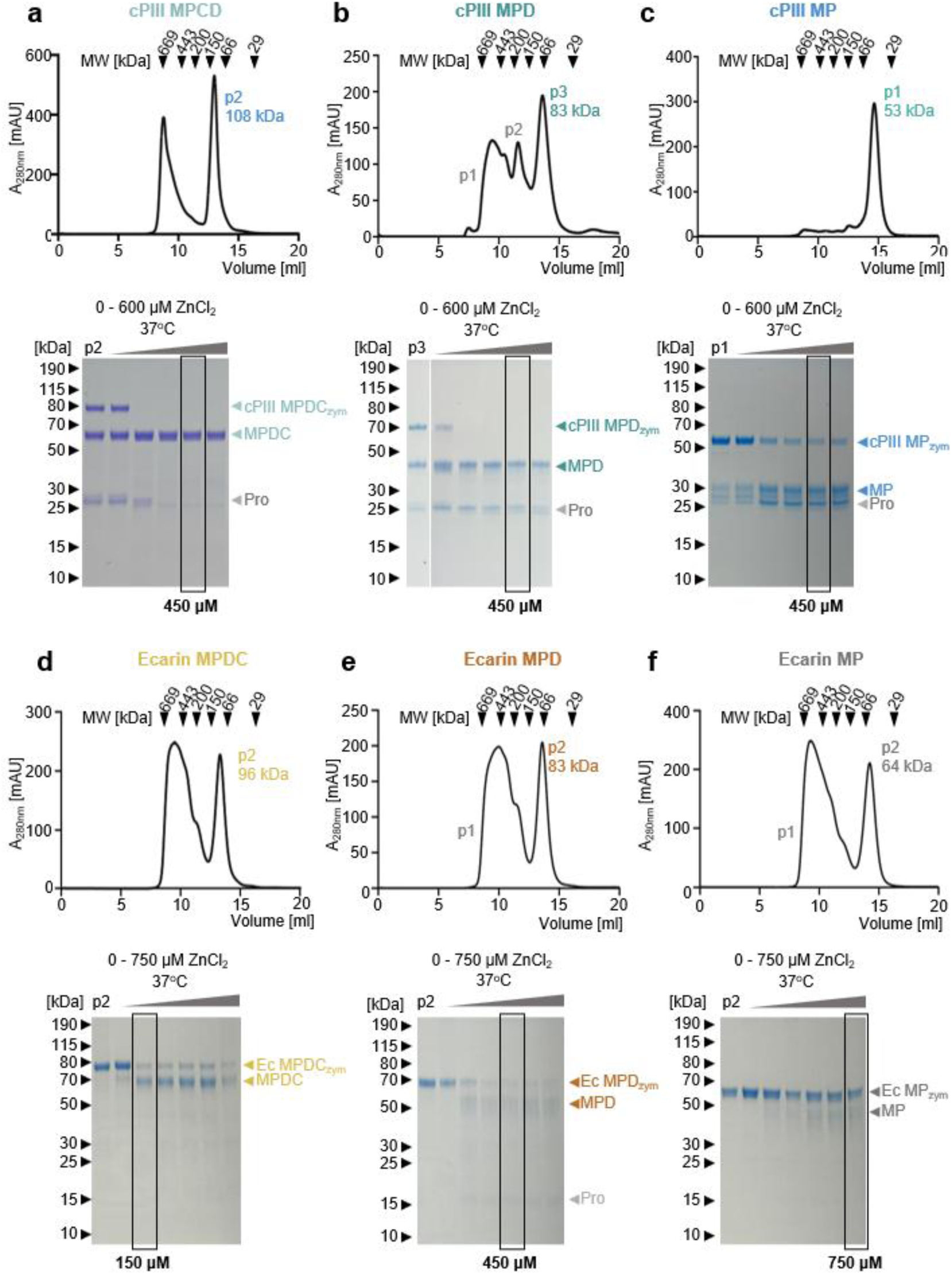
Purification and Activation of SVMP constructs. Size exclusion chromatograms of (**a**) cPIII MPDC, (**b**) cPIII MPD (**c**) cPIII MP, (**d**) Ecarin MPDC, (**e**) Ecarin MPD and (**f**) Ecarin MP SVMPs, following IMAC and IEX. Elution volumes of the MW calibration markers are indicated in the chromatograms as black arrows. SDS-PAGE gels show auto-activation of respective zymogens into mature enzymes, during 18hour incubations at increasing concentrations of ZnCl_2._(from 0 to 750 μM, in steps of 150 μM). cPIII MPDC and MPD were auto-activated at pH 8, cPIII MP and all Ecarin SVMP at pH 7. ZnCl_2_ concentrations used in activation experiments are boxed in black.

### Reduced prodomain degradation and auto-activation of cPIII Dis and C-rich domain deletion constructs

All three cPIII SVMPs (MPDC, MPD and MP) underwent partial activation during expression and purification, as previously reported for the full-size cPIII MPDC (Hall *et al*., 2026). Reducing SDS-PAGE indicated substantial prodomain cleavage, resulting in the mature form of the proteins. Despite this cleavage, the prodomain appears to remain associated with the metalloproteinase domain, as the protein eluted from SEC as a single prominent peak with an apparent molecular weight (MW) of ∼100 kDa (Figure 2a). The observed difference to the theoretical MW of the zymogen (70 kDa) is likely due to glycosylation (see below) (Hall *et al*., 2026). The yield of cPIII MPDC was 2.2 mg from 1 L Hi5 insect cell culture.

cPIII MPD (PII-like) displayed similar behaviour to full-length cPIII (Figure 2b), with partial cleavage of the prodomain from the mature protein. SEC resulted in a major peak eluting at ∼80 kDa (theoretical MW 59.6 kDa). A secondary peak of cPIII MPD was observed eluting at ∼11.5 mL, corresponding to dimeric protein (apparent MW ∼200 kDa. The presence of a void volume peak (∼9 mL) indicated some aggregation. The yield of monomeric cPIII MPD was 1.5 mg from 1 L Hi5 insect cell culture.

cPIII MP (PI-like) eluted in a single, well-defined peak at 14.5 mL corresponding to ∼53 kDa (theoretical MW: 47.6 kDa), with no significant aggregation detected (Figure 2c). cPIII MP also underwent partial cleavage of the prodomain, although a higher percentage of protein remained in the zymogen form than for cPIII MPD and MPDC. The yield obtained was 1.2 mg cPIII MP from 1 L Hi5 insect cell culture.

Full-length cPIII is predicted to comprise two potential glycosylation sites according to the NetNGlyc 1.0 server (Gupta & Brunak, 2002), with one site residing in the MP domain, and the other in the C-rich domain (Supplementary Figure 2a). To confirm glycosylation, the constructs were treated with Peptide N-glycosidase F (PNGase F) (Supplementary Figure 2b). Under reducing conditions, Coomassie-stained SDS-PAGE shows a clear downward shift of ∼5 kDa and ∼3 kDa for the bands corresponding to the PNGase F-treated cPIII MPDC mature protein and the cPIII MPD mature protein, respectively. The reduction in molecular weight of the cPIII MP zymogen after PNGase F treatment was less pronounced, consistent with the presence of one glycosylation site in the MP protein (Supplementary Figure 2b).

All three variants of cPIII zymogens could be auto-activated by addition of Zn^2+^ and overnight incubation at 37^°^C. Full-length cPIII MPDC was auto-activated by incubation with 150 μM ZnCl_2_ as shown by the disappearance of the zymogen band (∼77 kDa) on reducing SDS-PAGE, while at 450 μM Zn^2+^ concentration the prodomain was fully degraded (Figure 2a).

cPIII MPD (PII-like) was also auto-activated by 150 μM ZnCl_2_, however, its prodomain was not proteolysed, even in the presence of 600 μM ZnCl_2_ (Figure 2b). The cPIII MP (PI-like) showed inefficient auto-activation, with a significant amount of zymogen present after incubation even with 600 μM ZnCl_2_, and the prodomain remaining intact (Figure 2c).

For all subsequent SVMP activity assays, cPIII MPDC and MPD were activated by incubation with 450 μM ZnCl_2_. Because cPIII MP auto-activation was inefficient at pH 8, a pH screen was performed, indicating maximal but not complete activation at pH 6-7 (Supplementary Figure 3a). Therefore, cPIII MP was auto-activated by incubating with 750 μM Zn^2+^ in pH 7 buffer for all subsequent activity assays, in accordance with assay buffer requirements.

In conclusion, all three cPIII SVMP variants can be produced in milligram quantities by using baculovirus/insect cell expression. Protein yield decreased slightly with successive domain deletions. Auto-activation efficiency decreased with sequential domain removal. Full-length cPIII MPDC was most efficiently auto-activated and degraded its prodomain. cPIII MPD and MP failed to degrade their prodomains, and cPIII MP also failed to achieve complete auto-activation under the conditions tested.

### Deletion of Dis and C-rich domains from Ecarin impairs SVMP auto-activation efficiency

Ecarin variants (MPDC, MPD and MP) largely remained in their zymogen form during purification (Figure 2d-f). In contrast to the cPIII SVMPs (Figure 2a-c), they did not undergo complete spontaneous self-cleavage, but required addition of Zn^2+^ ions at 37^°^C for activation (Figure 2d-f). All three variants were expressed and purified in comparable amounts, with approximately 50% protein loss during SEC due to aggregation (elution at the column void volume, ∼9 ml). Monomeric Ecarin MPDC eluted with an observed MW of ∼96 kDa (Figure 2d), compared to the theoretical MW of 72.6 kDa, with the discrepancy likely attributed to glycosylation. Monomeric Ecarin MPD (PII-like) eluted at a MW of ∼83 kDa (Figure 2e), compared to the theoretical MW of 56 kDa, and monomeric Ecarin MP (PI-like) eluted with a MW of ∼64 kDa (Figure 2f), compared to the theoretical MW of 47 kDa. Production yields from 1 L Hi5 insect cell expressions were 2.4 mg for Ecarin MPDC, 2.7 mg for Ecarin MPD and 3.2 mg for Ecarin MP.

Full-length Ecarin is predicted to comprise four potential glycosylation sites according to the NetNGlyc 1.0 server (Supplementary Figure 2a) (Gupta & Brunak, 2002), with all four sites residing within the MP domain. The constructs were treated with PNGase F. A downward shift of ∼5 kDa was observed for all three bands corresponding to PNGase F-treated Ecarin constructs under reducing conditions on a Coomassie-stained SDS-PAGE gel (Supplementary Figure 2b), consistent with glycosylation of the Ecarin MP domain.

Partial auto-activation of Ecarin MPDC was observed with 150-750 μM ZnCl_2_ at pH 8, while Ecarin MPD and MP did not auto-activate under these conditions (data not shown). A subsequent pH screen revealed auto-activation was most pronounced at pH 6 – 7 (Supplementary Figure 3b–d). Subsequent ZnCl_2_ titrations for SVMP auto-activation (Figure 2d-f) were carried out in 50 mM HEPES, 150 mM NaCl, 2.5 mM CaCl_2_, pH 7 in accordance with assay buffer requirements. As observed for the cPIII domain deletion constructs (Figure 2a-c), progressive removal of the domains reduced the ability of Ecarin to undergo Zn^2+^-dependent auto-activation (Figure 2d-f). Ecarin MPDC was mostly auto-activated, accompanied by prodomain degradation (Figure 2d). Ecarin MPD (PII-like) could be almost completely auto-activated while the prodomain remained intact (Figure 2e). In contrast, Ecarin MP (PI-like) showed only partial auto-activation and no prodomain degradation, even at the highest concentrations of Zn^2+^ concentrations tested (Figure 2f). For all subsequent activity assays, Ecarin MPDC was activated at pH 7 with 150 μM ZnCl_2_, Ecarin MPD with 450 μM ZnCl_2_ and Ecarin MP with 750 μM ZnCl_2_.

Taken together, all Ecarin variants can be produced in milligram amounts (2.4 – 3.2 mg/L culture), and their auto-activation efficiency and prodomain degradation ability decrease progressively with sequential Dis and C-rich domain removal, consistent with our observations for cPIII SVMP variants.

### Deletion of cPIII Dis and C-rich domains reduces proteinase activity and cytotoxicity

Degradation of casein presents a general SVMP activity assay (Macêdo, 2016), whereas fibrinogen and prothrombin are substrates specific to certain SVMPs (Kini & Koh, 2016) and highlight differences in their haemotoxic activities. To assess the activity of our SVMP constructs, the zymogens were supplemented with optimal concentrations of ZnCl_2_ for overnight auto-activation (Figure 2). Subsequently, substrate and varying concentrations of enzyme were mixed and incubated at 37^°^C. Proteolysis was analysed by Coomassie-stained reducing SDS-PAGE (Figure 3a-c, Supplementary Figure 4). In all cases, indicated enzyme concentrations assume complete (100%) SVMP auto-activation by Zn^2+^.

**Figure 3.**
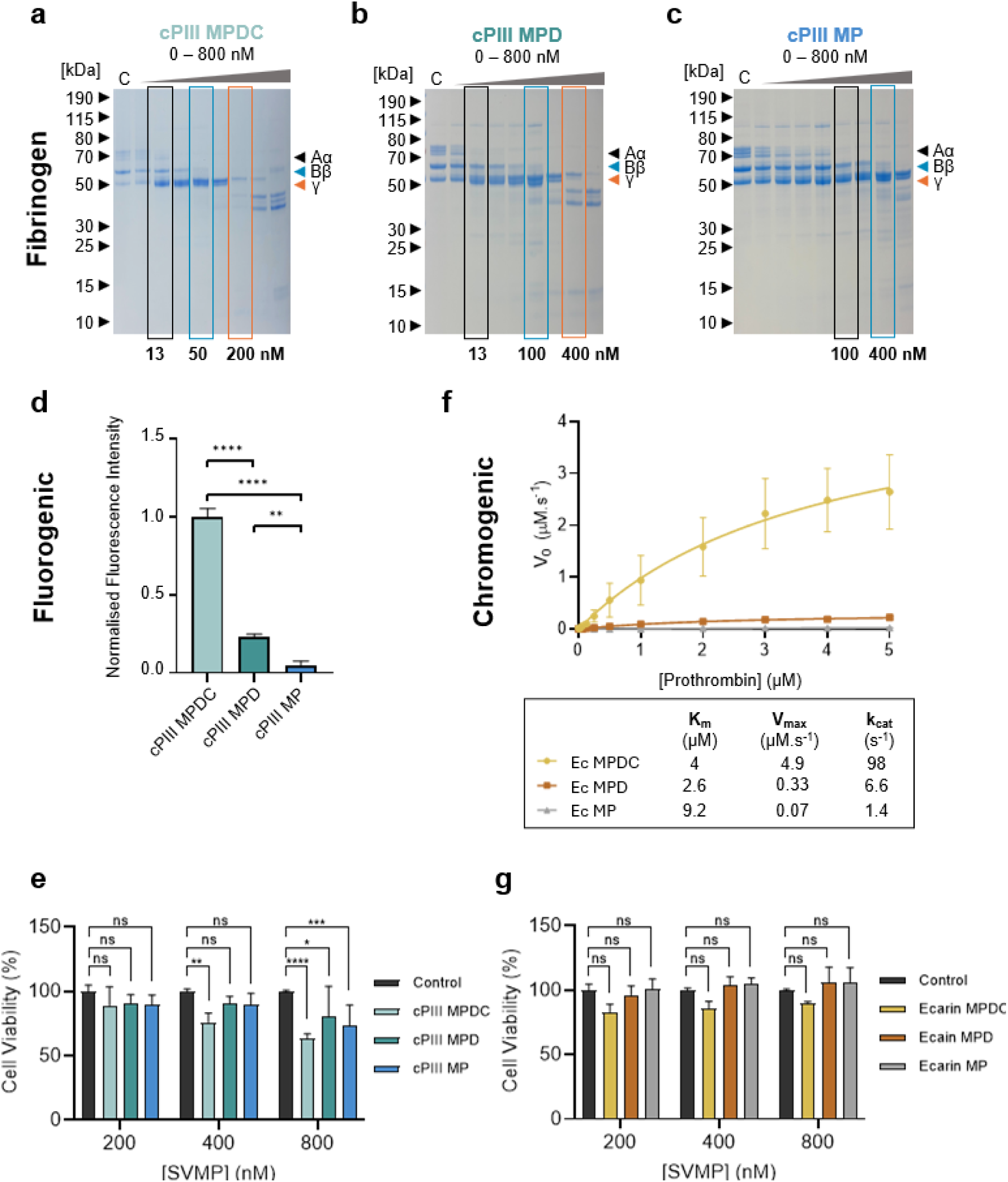
cPIII and Ecarin SVMP activity assays. Fibrinogen degradation following 18 hour incubation with as serial dilution of (**a**) cPIII MPDC, (**b**) cPIII MPD, and (**c**) cPIII MP. Orange arrowheads indicate the fibrinogen Aα, Bβ, and γ chains. (**d**) cPIII activity towards the fluorogenic peptide substrate ES010. (**e**) MTT cytotoxicity assay showing viability of HaCaT cells after 24-hour incubation with activated cPIII MPDC, MPD and MP SVMPs. (**f**) Michaelis–Menten analysis of prothrombin (PT) activation by Ecarin constructs. K_m_, V_max_ and k_cat_ values are tabulated below. (**g**) MTT cytotoxicity assay showing viability of HaCaT cells after 24-hour incubation with activated Ecarin MPDC, MPD and MP.

Casein was efficiently and completely degraded by cPIII, with clear proteinase activity exhibited at 100 nM for both cPIII MPDC and cPIII MPD (Supplementary Figure 4a,b). In contrast, cPIII MP (PI-like) required four times as much enzyme (400 nM) to reach a comparable effect (Supplementary Figure 4c).

Fibrinogen degradation by cPIII was reported previously (Hall *et al*., 2026). Full-length cPIII MPDC and cPIII MPD (PII-like) were most active, causing preferential degradation of the Aα chain at very low concentrations (13 nM), followed by the degradation of the Bβ chain at higher concentrations (50 nM MPDC and 100 nM MPD) (Figure 3a, b). Notably, degradation of the γ chain was observed at 200 nM MPDC and 400 nM MPD (Figure 3a,b). In contrast, cPIII MP (PI-like) exhibited the lowest activity, with detectable Aα chain degradation activity at 100 nM, Bβ chain degradation activity at 400 nM and minimal γ chain cleavage even at 800 nM (Figure 3c).

ES010, a PIII-specific fluorogenic peptide substrate, is efficiently cleaved by full-length cPIII (Hall *et al*., 2026). To test ES010 cleavage, 100 nM SVMP zymogen (corresponding to approximately 700 ng cPIII MPDC, 600 ng MPD and 475 ng MP) was used. Fluorescence at the start of the experiment was subtracted from the 15-minute endpoint and normalised to the mean value for cPIII MPDC (Figure 3d). Full-length MPDC activity was significantly higher than that of MPD and MP for this substrate. Moreover, MPD (PII-like) activity was significantly higher than MP (PI-like) activity (*p* = 0.0018). Thus, successive domain deletion from cPIII resulted in a progressive reduction in activity against the fluorogenic peptide substrate in our experiments.

We analysed the activity of cPIII constructs towards prothrombin (Supplementary Figure 5). No thrombin generation was detected in our experiments, however, pronounced degradation bands were observed after overnight incubation with full-length MPDC and MPD (PII-like) (Supplementary Figure 5b-d). Prothrombin activation to thrombin was not observed, nor did any cPIII construct display activity toward the chromogenic substrate S-2238 (Supplementary Figure 6a), which is used to measure thrombin generation. Statistical analysis showed that all cPIII constructs had non-significant activity when compared to a no SVMP control. Therefore, the cPIII constructs are not prothrombin-activating proteins.

Cytotoxic effects of cPIII constructs were assessed using an MTT viability assay in human umbilical vein endothelial cells (HUVEC) following a 24 hour exposure to activated enzymes (Figure 3e). At 200 nM, none of the constructs induced a significant reduction in cell viability relative to the control. At 400 nM, only the full-length cPIII MPDC showed a statistically significant reduction in viability. At 800 nM, all cPIII constructs reduced cell viability significantly, MPDC exhibited the strongest cytotoxic effect, followed by MPD, while the most truncated construct (MP) showed the weakest effect on cell viability, confirming that deletion of Dis and C-rich domains reduces both proteinase activity and cytotoxicity.

Taken together, the Dis and C-rich domain deletion constructs of cPIII showed no changes in their substrate spectrum and sequentially reduced activity compared to the full-length MPDC construct.

### N-glycosylation modulates proteinase activity of cPIII constructs

To assess the contribution of N-linked glycosylation to enzymatic function, cPIII constructs were treated overnight with PNGase F under non-reducing conditions (Supplementary Figure 2b). ES010 cleavage was then compared between the glycosylated and deglycosylated forms of the cPIII constructs (Supplementary Figure 2c). Deglycosylation resulted in construct-dependent effects on catalytic activity. For full-length cPIII MPDC and truncated cPIII MP, removal of N-linked glycans significantly reduced enzymatic activity, indicating a role for glycosylation in maintaining catalytic function. Surprisingly, cPIII MPD (PII-like) displayed an increase in activity following deglycosylation, suggesting a potentially inhibitory effect of glycosylation in this truncated construct.

In summary, glycosylation exerts construct-dependent effects on cPIII SVMP activity, enhancing catalytic efficiency in full-length and MP constructs, while reducing activity in the MPD (PII-like) construct, suggesting that N-linked glycans modulate enzymatic function in a structure-dependent manner.

### Deletion of Dis and C-rich domains of Ecarin leads to reduced proteolytic activity, prothrombin activation and blood clotting

Ecarin displays much narrower specificity compared to the broad substrate profile of cPIII (Paine & Laing, 2004). Ecarin MPDC was able to degrade casein, albeit at higher concentrations as compared to cPIII MPDC (300 nM and 100 nM, respectively) (Supplementary Figure 4d). This activity was characterised by the appearance of discrete lower molecular weight fragments rather than the near-complete substrate digestion observed for cPIII MPDC. Casein cleavage by Ecarin MPDC was detected at 300 nM, whereas Ecarin MPD required 1200 nM for detectable activity (Supplementary Figure 4e). Ecarin MP was not active at the conditions tested (Supplementary Figure 4f).

Ecarin is known for its highly specific activation of prothrombin to thrombin (Paine & Laing, 2004). Consistent with this, Ecarin MPDC efficiently generated thrombin at concentrations as low as 13 nM following 1 hour incubation (Figure 4a). Successive domain deletion resulted in a marked reduction in activity, with Ecarin MPD (PII-like) showing only partial activity at 200 nM and Ecarin MP (PI-like) showing minimal to no activity. With Ecarin MPDC and MPD, thrombin formation (∼31 kDa under reducing conditions) was confirmed by its characteristic upward shift to ∼36 kDa in non-reducing SDS-PAGE (Supplementary Figure 5e), reflecting retention of disulphide bonds between the thrombin A and B chains resulting in a higher apparent molecular weight under non-reducing conditions (Krishnaswamy *et al*, 1987). No detectable activity by Ecarin MP occurred within a 1 hour incubation period, and while some activity was seen after 18 hours, a detectable thrombin band was not observed (Supplementary Figure 5g), indicating Ecarin MP was degrading prothrombin rather than liberating thrombin.

**Figure 4:**
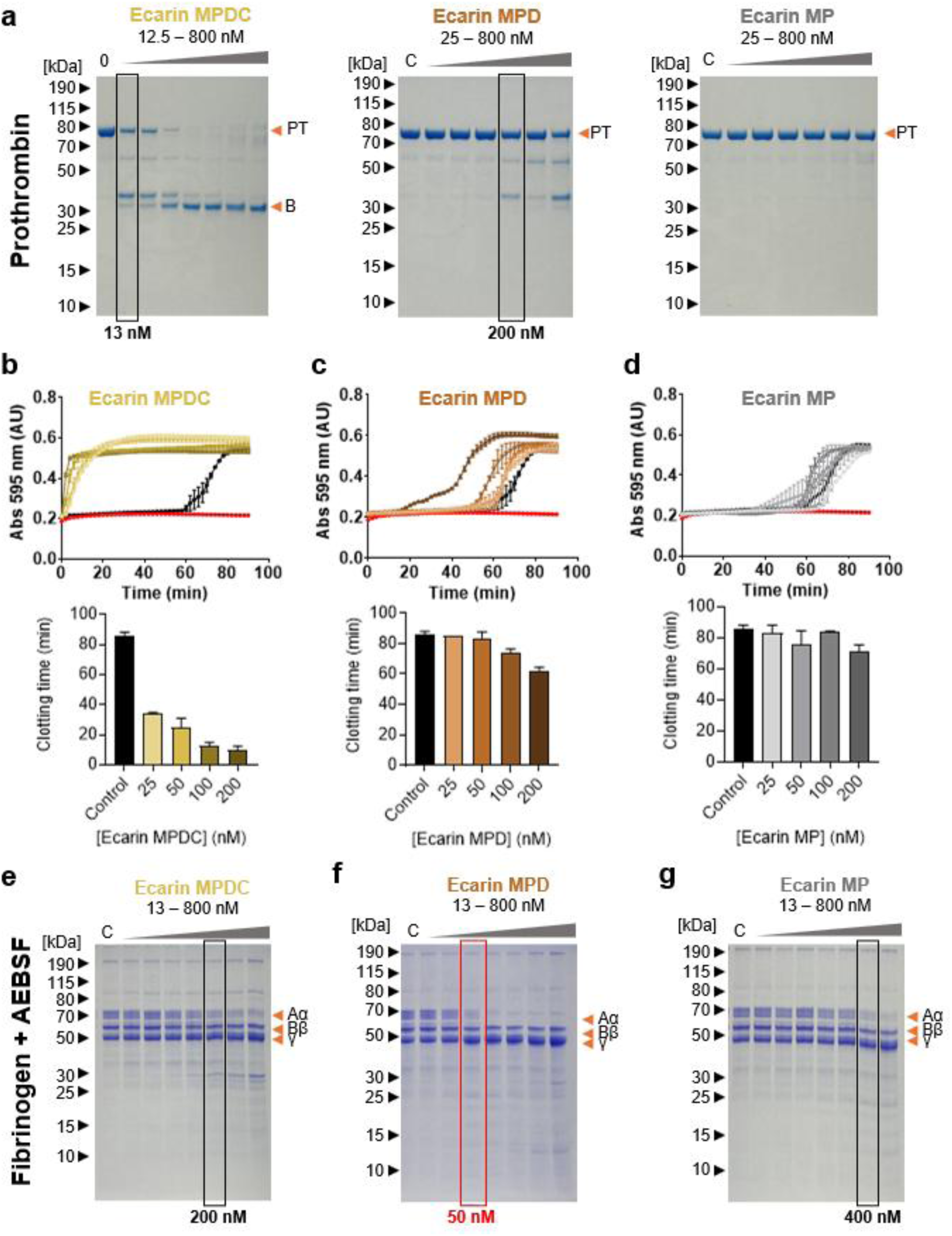
Ecarin activity assays. (**a**) SDS-PAGE analysis of prothrombin activation following 1 h incubation with a serial dilution of Ecarin MPDC, MPD, and MP. Orange arrowheads indicate prothrombin and the B-chain of α-thrombin. C, control: incubation in the presence of 20 mM EDTA. (**b**-**d**) Plasma clotting assay with (**b**) Ecarin MPDC, (**c**) Ecarin MPD and (**d**) Ecarin MP in the presence of calcium. Clotting time was determined by the time required to reach a stable absorbance plateau, corresponding to maximal clot formation. (**e**-**g**) Fibrinogen degradation in the presence of the serine-protease inhibitor, AEBSF, with a serial dilution of (**e**) Ecarin MPDC, (**f**) Ecarin MPD and (**g**) Ecarin MP. Orange arrowheads: fibrinogen chains Aα, Bβ and γ. C (control): pre-incubation of SVMP with 20 mM EDTA. Boxes highlight detectable degradation.

Kinetic analysis of cleavage of the S-2238 chromogenic substrate, which is indicative of thrombin formation, further supported activation of prothrombin by Ecarin rather than prothrombin degradation. Ecarin MPDC exhibited strong activity with an estimated K_M_ of 4 μM and k_cat_ of 98 s⁻¹ resulting in a k_cat_ / K_M_ value of 2.45×10^7^ M^−1^s^−1^ (Figure 3f), comparable to previously reported values for recombinant Ecarin (K_M_ ∼4.4 μM) (Misson Mindrebo *et al*., 2024). Activity was strongly reduced for truncated Ecarin constructs (Figure 3f).

### Ecarin activity is enhanced by N-glycosylation

Next, we investigated the impact of glycosylation of Ecarin on its activity. *In silico* analysis identified four putative N-linked glycosylation sites in Ecarin, all located within the metalloproteinase (MP) domain (Supplementary Figure 2a). In agreement, treatment with PNGase F produced a consistent downward shift in SDS-PAGE banding patterns across all three constructs, confirming the presence of N-linked glycans in the MP domain (Supplementary Figure 2b). We tested the de-glycosylated Ecarin constructs for cleavage of the S-2238 chromogenic substrate. To exclude effects of PNGase F on assay performance, a PNGase F-only control was included in the S-2238 assay, showing no significant difference from a control reaction without any enzyme added (data not shown), confirming that PNGase F does not affect substrate turnover in these assay conditions. Deglycosylation resulted in a significant reduction in S-2238 cleavage for both Ecarin MPDC and MPD (Supplementary Figure 2d). No significant difference was observed between glycosylated and deglycosylated Ecarin MP; both were catalytically virtually inactive (Supplementary Figure 2e).

### Ecarin’s Dis and C-rich domains are required for efficient plasma clotting

Thrombin activation by Ecarin leads to plasma clot formation. The three Ecarin constructs were assessed for their ability to induce plasma clot formation using standard turbidity-based clotting assays (Larsen & Hvas, 2020). Ecarin was reported to be a Ca²⁺-independent prothrombin activator (Kornalik & Blomback, 1975), nonetheless, we included Ca^2+^ in our buffers for consistency. Ca^2+^-containing buffer (no Ecarin added) induced plasma clotting after 84 min in our assay (Figure 4). In comparison, Ecarin MPDC induced rapid clot formation, with clotting observed within 8 min at 200 nM (Figure 4b). Under the same conditions, Ecarin MPD (PII-like) required 65 min, while Ecarin MP (PI-like) required 76 min (Figure 4c,d).

### Deletion of the C-rich domain of Ecarin leads to increased fibrinogen degradation

Fibrinogen is not a reported substrate of Ecarin (Jonebring *et al*, 2012; Paine & Laing, 2004). Fibrinogen degradation assays were performed in the presence of the serine protease inhibitor AEBSF (Figure 4e-g). Ecarin MPDC and MP (PI-like) showed minimal fibrinogen Aα chain degradation at 200–400 nM (Figure 4e,g). We note that the auto-activation of MP is incomplete (c.f. Figure 2f), and remaining zymogen in the mixture makes it difficult to estimate the effective enzyme concentration. In contrast, Ecarin MPD (PII-like) detectably degraded the Aα chain at 50 nM (Figure 4f), suggesting that the C-rich domain of Ecarin may restrict access of fibrinogen substrates to the MP active site.

### Insect cell-produced, recombinant Ecarin closely recapitulates the enzymatic activity of native Ecarin

We asked if insect cell-produced recombinant Ecarin exhibits the expected biochemical properties of native Ecarin. We compared our recombinant construct to a pooled native Ecarin preparation, purified from *Echis ocellatus* and *Echis romani*. We could not perform the same comparison for cPIII due to lack of native cPIII purified in sufficient quality and quantity from venom.

Native Ecarin pool migrated as a set of three major bands between ∼60–70 kDa on reducing SDS-PAGE, with the lowest band corresponding to the expected MW of Ecarin at 56 kDa (Figure 5a). Recombinant Ecarin was expressed as a zymogen with a band corresponding to ∼80 kDa in reducing SDS-PAGE. Following Zn²⁺-dependent auto-activation at 37°C for 18 hours, a shift to <70 kDa is observed, consistent with cleavage of the prodomain and activation of the recombinant protein (Figure 5a). As expected, a corresponding shift was not observed for native Ecarin purified from venom, which comprises the mature active SVMP lacking the prodomain..

**Figure 5.**
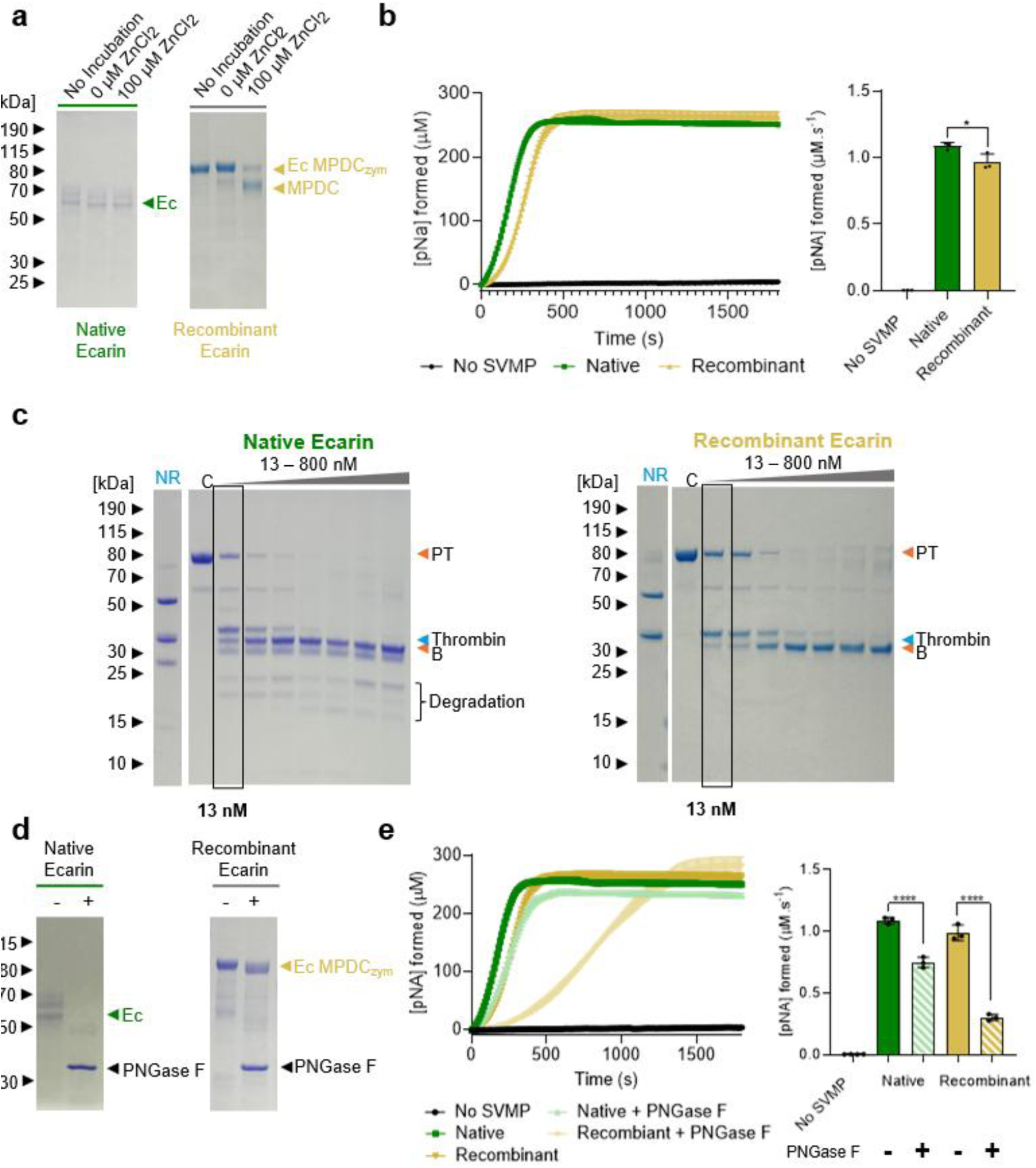
Comparison of native and recombinant Ecarin activity. (**a**) SDS-PAGE analysis of native and recombinant Ecarin preparations. Comparison of prothrombin (PT) activation by native and recombinant Ecarin (MPDC) using the (**b**) chromogenic substrate S-2238 and (**c**) SDS-PAGE analysis following 1 hour incubation. Orange arrowheads indicate prothrombin and the B-chain of α-thrombin, while blue arrowheads indicate α-thrombin in non-reducing (NR) lanes. Removal of N-glycans by PNGase F treatment assessed by (**d**) SDS-PAGE and (**e**) comparison of PT activation by native and recombinant deglycosylated Ecarin.

Prothrombin activation was assessed using the chromogenic substrate S-2238. Both native and recombinant Ecarin exhibited similar prothrombinase activity at 100 nM, demonstrating that recombinant Ecarin is capable of reproducing the expected functional activity of the native toxin (Figure 5b). A minor, but statistically significant, difference was detected (Figure 5b), which may reflect sequence variation between species (*E. ocellatus* and *E. romani vs. E. pyramidum leakeyi*), or incomplete activation of the recombinant zymogen, as evidenced by the residual zymogen band observed in SDS-PAGE following activation (Figure 2d, 5a).

SDS-PAGE analysis of prothrombin activation further demonstrated comparable conversion of prothrombin to thrombin by the recombinant preparation (Figure 5c). Similar enzyme concentrations (13 nM) were required to achieve thrombin formation in the same time period (1 hour). The native Ecarin pool generated additional lower MW bands, consistent with alternate proteolytic degradation activities or prothrombin degradation products, which were not observed in recombinant preparations, indicating an absence of contaminating proteinases and thus a higher purity of the recombinant Ecarin.

Treatment with PNGase F resulted in a reduction in molecular weight for both native and recombinant Ecarin (Figure 5d). The magnitude of this shift was more pronounced in the native preparation, indicative of differences in the glycosylation patterns between the native and insect-cell-produced proteins. PNGase F removes N-linked sugars, the predominant form of glycosylation present in native SVMPs (Fox & Serrano, 2005; Tarentino *et al*, 1985). N-linked glycosylation in snakes closely resembles the glycosylation of mammalian proteins (Andrade-Silva *et al*, 2018; Soares & Oliveira, 2009), while recombinant SVMPs produced in baculovirus-infected insect cells are expected to carry less complex, predominantly paucimannosidic insect-type glycans (Shi & Jarvis, 2007). Nonetheless, functional assessment following deglycosylation demonstrated a significant reduction in S-2238 activity for both native and recombinant Ecarin (Figure 5e), conveying that N-linked glycosylation contributes to catalytic efficiency in both protein preparations.

Overall, these results demonstrate that the insect cell-produced Ecarin exhibits the expected activity profile of its snake-produced counterpart, highlighting the utility of the baculovirus/insect cell expression system to recombinantly produce catalytically active authentic SVMPs with native-like properties in quantities and quality for functional and mechanistic dissection.

## DISCUSSION

Snake venom metalloproteinases (SVMPs) represent a striking example of protein family diversification driven by gene duplication and lineage-specific domain loss (Casewell *et al*, 2012; Giorgianni *et al*, 2020; Sanz & Calvete, 2016). These evolutionary processes have generated a structurally diverse set of toxins ranging from the simple PI SVMPs, consisting solely of a metalloproteinase domain, to the multidomain PIII SVMPs that additionally contain Dis and C-rich accessory domains. Although the evolutionary origin of this modular architecture is well established (Casewell *et al*., 2011; Giorgianni *et al*., 2020; Sanz & Calvete, 2016), the mechanistic contribution of the accessory domains to substrate recognition and catalysis remains incompletely understood. In this study, we addressed this question using systematic domain-deletion variants of two functionally distinct PIII SVMPs.

Within PIII SVMPs, the accessory domains form a compact arrangement adjacent to the catalytic MP domain (Figure 1), suggesting potential roles in regulating substrate access and positioning. Based on this architecture, we considered two not mutually exclusive models: (i) a ‘recruitment-feeding’ role in which PIII SVMPs facilitate productive substrate engagement and orientation for catalysis, and (ii) a steric ‘gatekeeping’ role in which the accessory domains restrict substrate access to the MP catalytic cleft. By using deletion constructs, we dissected here the relative contribution of the Dis and C-rich domains to enzymatic activity and substrate specificity.

Recently, we reported successful recombinant production of cytotoxic and haemotoxic SVMPs using the MultiBac baculovirus/insect cell system, which enabled systematic functional interrogation of toxins which were previously not accessible (Hall *et al*., 2026). Here, we used this expression platform to generate full-length SVMPs and domain truncation constructs. We dissected the contributions of the Dis and C-rich domains in two functionally divergent PIII SVMPs: a broadly toxic enzyme (cPIII) that we have previously characterised (Hall *et al*., 2026), and the highly procoagulant, well-characterised thrombin activator Ecarin (Kini & Koh, 2016; Nishida *et al*., 1995). Both proteins and their corresponding truncation variants were successfully expressed as zymogens and activated by incubation with Zn²⁺ (Figure 2). Full-length proteins showed efficient auto-activation and complete prodomain degradation, whereas deletion of Dis and C-rich domains prevented the degradation of the prodomain. Moreover, the efficiency of auto-activation decreased with stepwise deletion of domains.

Activity assays showed that in cPIII SVMP, sequential domain deletion results in a consistent reduction in catalytic activity across multiple substrate classes (Figure 3). The MP domain alone retained measurable activity, but inclusion of the Dis and C-rich domain progressively enhanced catalytic efficiency, particularly toward the ES010 fluorogenic substrate. ES010 comprises a central PLGL cleavage motif, derived from sequences preferred by many matrix metalloproteinases and some ADAM proteinases which are evolutionarily related to SVMPs (Takeda, 2016). The PLGL motif in ES010 resembles motifs present in several extracellular matrix proteins that these proteinases target, including collagen-derived substrates (Neumann *et al*, 2004).

Our cPIII findings do not support a simple gatekeeper model. Instead, they are more consistent with a model in which accessory domains enhance catalysis by promoting substrate engagement and productive alignment with the active site (recruitment-feeding). The pronounced effect on small peptide substrates suggests that the C-rich domain is particularly important in stabilising low-affinity interactions, whereas larger protein substrates may engage additional surface contacts with the MP and Dis domain, partially compensating for C-rich domain loss. The strongly reduced activity of the cPIII deletion constructs is in agreement with our previous finding that PII and PI SVMPs did not exhibit any substantial cleavage of ES010 (Hall *et al*., 2026).

Ecarin displays a highly restricted substrate profile dominated by prothrombin activation (Kornalik & Blomback, 1975). In good agreement, full-length Ecarin (MPDC) efficiently converted prothrombin to thrombin at concentrations as low as 13 nM, whereas sequential deletion of accessory domains caused a marked loss of activity (Figure 4). The C-rich domain is particularly important for efficient prothrombin activation, with its removal resulting in strongly reduced catalytic efficiency. This was confirmed by plasma clotting assays (Figure 4b-d). In summary, these data support a model in which the C-rich domain contributes to substrate recognition and productive positioning of prothrombin within the MP catalytic cleft.

Removal of Ecarin’s C-rich domain did not result in a general increase in catalytic activity across all substrates tested. No enhancement was observed for prothrombin cleavage, casein degradation, or S-2238 cleavage upon deletion of this domain (Figure 3,4). However, Ecarin MPD displayed increased fibrinogenolytic activity relative to MPDC (Figure 4e,f). This indicates that the C-rich domain does function as a ‘gatekeeper’ for this substrate. Thus, the C-rich domain could restrict access or prevent productive engagement with specific protein substrates such as fibrinogen.

We also analysed the importance of N-linked glycosylation of these SVMPs (Supplementary Figure 2). Computational analysis revealed multiple modification sites within full-length cPIII and Ecarin and their truncation variants. Deglycosylation with PNGase F resulted in a shift to lower molecular weight in all protein constructs, confirming successful removal of N-linked glycans. Functionally, glycosylation clearly contributed to catalytic activity (Supplementary Figure 2). Deglycosylation significantly reduced prothrombin-activating activity in full-length MPDC and MPD Ecarin constructs. The Ecarin MP domain displayed virtually no activity, and thus no reduction in activity due to deglycosylation could be observed. Of note, deglycosylation of the cPIII MPD construct resulted in increased activity, while the full-length MPDC and MP constructs showed reduced activity upon removal of the sugars. We hypothesise that the increased activity of cPIII MPD after PNGase treatment could be due to a more open or exposed active site, or facilitated substrate recruitment, in the deglycosylated protein. Taken together, glycosylation of these SVMPs modulates enzyme efficiency, potentially by influencing structural stability or local conformational dynamics.

We further assessed the biochemical properties of recombinant Ecarin using a native Ecarin preparation as a reference. Although direct comparison between the two is limited by differences in protein source, amino acid sequence and purity, both preparations exhibited the characteristic prothrombin-activating activity, with comparable concentration dependence and thrombin generation profiles. Both recombinant and native Ecarin generated thrombin, evidenced by the ∼30 kDa band under reducing conditions, and the ∼36 kDa disulphide-linked form under non-reducing conditions. The additional lower MW species observed for native Ecarin is likely caused by contaminating proteinases from the venom rather than differences in the Ecarin-mediated activation mechanism. We conclude that the baculovirus/insect cell expression system provides a robust platform suitable for producing catalytically active native-like SVMPs.

Our findings are consistent with previous studies showing that PIII SVMPs are more haemotoxic than SVMP PIs, which comprise the MP domain only (Gutierrez *et al*, 2016). Our data also support a unified model in which PIII SVMP accessory domains act primarily as modulators of substrate positioning and catalytic efficiency rather than absolute determinants of substrate access. In cPIII, the Dis and C-rich domains together enhance catalytic efficiency across multiple substrate classes by enhancing productive engagement with diverse targets. In Ecarin, the C-rich domain is particularly important to enable highly efficient and selective prothrombin activation. We conclude that rather than functioning strictly as ‘gatekeepers’ or ‘substrate feeders’, the Dis and C-rich domains tune the balance between general accessibility of the MP active site and productive engagement with specific substrates.

In which ways could these findings inform rational antivenom development? Neutralising SVMPs remains a major challenge because these toxins are principal mediators of local tissue destruction, haemorrhage and coagulopathy following viper envenomation (Bittenbinder *et al*., 2024). Our study indicates that although catalytic activity resides within the MP domain, efficient substrate recognition and productive catalysis depend heavily on the Dis and C-rich accessory domains. In principle, direct inhibition of the catalytic site would provide the most effective means of abolishing proteolytic activity. However, the catalytic cleft is highly conserved among metalloproteinases, making it difficult to generate selective binders without cross-reactivity toward endogenous metalloproteinases. Furthermore, direct inhibition of the catalytic site is particularly challenging for antibody-based therapeutics. Antibodies would need to mimic substrate interactions to achieve effective inhibition and could themselves become susceptible to proteolytic cleavage. In contrast, the accessory domains are structurally diverse, solvent exposed, and likely mediate substrate recruitment through distinct substrate-binding surfaces that are separate from the catalytic cleft. Our data show that deletion of these domains markedly reduces enzymatic activity, suggesting that disruption of substrate recruitment, and potentially steric obstruction of active site access, may provide an effective alternative mechanism for SVMP neutralisation.

Several studies support the Dis and C-rich domains as promising therapeutic epitopes for recombinant antivenom development. Recent advances in recombinant antibody discovery have generated both monoclonal antibodies and nanobody formats capable of neutralising snake venom toxins (Ahmadi *et al*, 2025; Chavanayarn *et al*, 2012; Prado *et al*, 2016; Richard *et al*, 2013). Although relatively few studies have mapped antibody epitopes on SVMPs, emerging structural and functional evidence indicates that the Dis and C-rich accessory domains constitute effective targets for neutralisation. A notable example is the recently described synthetic neutralising antibody against Ecarin, which was shown by structural analysis to bind predominantly to the C-rich domain (Misson Mindrebo *et al*., 2024). Rather than occluding the catalytic zinc-binding site, the antibody prevented productive engagement of prothrombin, thereby inhibiting toxin activity through disruption of substrate recognition and positioning (Misson Mindrebo *et al*., 2024). In a similar vein, polyclonal antibodies generated against the Dis and C-rich domains of *Echis ocellatus* SVMPs significantly reduced haemorrhagic activity both *in vitro* and *in vivo*, demonstrating that these accessory domains contain functionally important neutralising epitopes (Hasson, 2017). Consistent with this, the monoclonal antibody MAJar3 recognises an epitope within the Dis region of the PIII SVMP jararhagin and effectively neutralises its haemorrhagic activity (Tanjoni *et al*, 2003).

Evidence from the closely related ADAM family further supports this strategy. Multiple neutralising antibodies have been developed against the C-rich domains of ADAM proteases, where they inhibit substrate recognition or receptor interactions without directly targeting the catalytic active site. For example, monoclonal antibodies against the C-rich domain of ADAM10 inhibit ephrin cleavage by disrupting substrate recognition, while a recently described antibody against the C-rich domain of ADAM17 blocks substrate processing through interference with substrate-interacting surfaces (Atapattu *et al*, 2012; Saha *et al*, 2024). Recombinant expression of isolated accessory domains has also enabled the generation of domain-specific antibodies, demonstrating that these regions are immunogenic and readily accessible for antibody binding (Atapattu *et al*., 2012; Saha *et al*., 2024). Given the high structural conservation between ADAM proteins and PIII SVMPs, these findings reinforce the concept that accessory-domain-directed biologics, including monoclonal antibodies and nanobodies, represent a feasible and potentially highly selective strategy for next-generation antivenom development.

## MATERIALS AND METHODS

### SVMP expression construct design

Ecarin (UniProt ID: Q90495) and the cytotoxic PIII (cPIII, UniProt ID: E9JG34 (Hall *et al*., 2026)) were designed by fusing an Avi- and His-tag (GGSGLNDIFEAQKIEWHEHHHHHHHH*) to the C terminus of the native zymogen sequence. The N terminus of the zymogen retained the native snake signal sequence immediately upstream of the prodomain. Sequences were codon-optimised for *Spodoptera frugiperda*, synthesised (Genscript) and inserted into the pACEBac1 plasmid (Geneva Biotech), which served as the donor vector in the MultiBac system (Sari *et al*, 2016).

Full-length proteins are described here as Ecarin MPDC and cPIII MPDC. Domain deletion constructs were designed by removing the C-rich (Ecarin MPD, cPIII MPD), or both the Dis and C-rich domains (Ecarin MP, cPIII MP). Domain boundaries were identified using the UniProt entries (Ecarin: Q90495, cPIII: E9JG34) and domain deletion constructs were designed accordingly: cPIII MPD: residues 1-493, cPIII MP: residues 1-406, Ecarin MPD: residues 1-491, Ecarin MP: residues 1-404 (with position 1 corresponding to the start of the protein with signal sequence). In all cases, the Avi- and His-tags were fused to the C terminus, separated by a GGS linker.

### Recombinant SVMP expression and purification

Each SVMP-containing pACEBac1 plasmid was expressed using the MultiBac baculovirus/insect cell expression system as previously described (Hall *et al*., 2026). Briefly, SVMPs were expressed in Hi5 insect cells at 19°C, in ESF 921 Insect Cell Culture Medium (Expression Systems). Expression was monitored by measuring the YFP fluorescence (encoded in the baculoviral genome) and cell viability (by determining the percentage of live cells using Trypan Blue). Cultures were harvested 5 days post-infection or when cell viability decreased below 80%, whichever occurred sooner. After harvesting the media, secreted SVMPs were immediately purified following the three-step purification protocol as outlined previously (Hall *et al*., 2026), except that all buffers were supplemented with 2.5 mM CaCl_2_.

### SVMP enzyme activation

SVMPs were activated by incubation with 0 – 750 μM ZnCl_2_ at 37°C for 18 hours as described previously (Hall *et al*., 2026). Activation was assessed by Coomassie-stained SDS-PAGE, monitoring the disappearance of the zymogen band.

For SVMPs that were not efficiently activated at pH 8, a pH titration screen was performed using 50 mM buffers at the indicated pH values (pH 5: sodium acetate; pH 6: MES; pH 7: HEPES; pH 8 [at 37°C]: Tris-HCl; pH 9-10: glycine-NaOH), supplemented with 150 mM NaCl, 2.5 mM CaCl_2_ and 450 μM ZnCl_2_.

### Casein, fibrinogen and prothrombin degradation assays

Casein and fibrinogen degradation were tested as examples for general and specific substrates, respectively (Hall *et al*., 2026). Briefly, SVMPs were first activated with Zn^2+^ as described above. Then, 0-1.2 μM activated SVMP (based on zymogen concentration prior to activation) was incubated for 18 hours at 37°C with casein or fibrinogen at a final concentration of 0.525 mg/mL (1.5 μM casein, 23 μM fibrinogen) in a reaction volume of 20 μL. For Ecarin MPDC, MPD and MP, prothrombin was used as the SVMP specific substrate. Ecarin-based SVMPs (0-1.2 μM) were incubated with a final concentration of 0.36 mg/mL prothrombin (2.5 μM) in a reaction volume of 20 μL for 1 hour (MPDC) or 18 hours (MPD, MP) at 37°C. The negative control included the addition of 5 mM EDTA as a metal chelator and SVMP inhibitor. Reactions were stopped with the addition of Protein Gel Loading Dye and heated to 95°C.

Substrate degradation or thrombin activation was analysed by Coomassie stained reducing SDS-PAGE gels. Thrombin generation was further confirmed by non-reducing SDS-PAGE, whereby the ∼30 kDa band shifts upwards to ∼36 kDa, due to the A and B chains remaining linked by disulphide bonds.

### Fluorogenic peptide substrate cleavage assay

Enzymes were activated with Zn^2+^ as described above and then diluted in reaction buffer (50 mM Tris-HCl, 150 mM NaCl, 2.5 mM CaCl_2_, pH 8). Subsequently, 700 ng cPIII MPDC, 600 ng cPIII MPD or 475 ng cPIII MP enzyme were added to each well, resulting in a final concentration of 100 nM based on the SVMP zymogen concentration at the start of the experiment. ES010 substrate (R&D Systems Inc.) was diluted in reaction buffer and used at a final concentration of 10 μM in a total reaction volume of 100 μL in a black 96 well plate (Thermo Fisher Scientific). Fluorescence was followed for 1 hour at 25°C using a BioTek Synergy Neo2 instrument, with an excitation wavelength of 320 nm and an emission wavelength of 405 nm. Readings were taken in triplicate and normalised to the mean value for cPIII MPDC. Data were plotted using GraphPad (Prism), and statistical significance was assessed by one-way ANOVA followed by Tukey’s multiple comparison tests.

### Chromogenic peptide substrate cleavage assay

S-2238 substrate (hydrochloride) (Cayman Chemical) was used as a chromogenic substrate to kinetically measure Ecarin-mediated activation of prothrombin. Ecarin converts prothrombin to thrombin, which cleaves the terminal Arg-pNA bond in S-2238 to release p-nitroaniline (pNA), measurable at 405 nm (Johnson *et al*, 2020; Jonebring *et al*., 2012).

SVMPs were incubated with Zn^2+^ as described above and then pre-heated to 37°C prior to the reaction. For reactions used to calculate apparent Km and kcat values, Ecarin construct concentrations of 50 nM, based on the MW of the zymogen, were prepared in 96-well clear microplate (Greiner Bio-One) before the addition of 0.5 mM S-2238 and varying concentrations of human prothrombin (0-5 µM) in reaction buffer (50 mM HEPES, 150 mM NaCl, 2.5 mM CaCl_2_, pH 8). Absorbance at 405 nm was recorded in 10 second intervals for 30 minutes, at 37°C using a BioTek Synergy Neo2 instrument, with reactions being run in triplicate, corrected for baseline activity by subtracting t=0 and converted to concentration of pNA formed using a calibration curve generated from endpoint absorbance values (Supplementary Figure 7). Initial velocities were calculated and fitted to the Michaelis-Menten equation using GraphPad (Prism).

For additional assays, reactions were conducted in clear, flat-bottomed half-area 96-well microplates (Greiner Bio-One), and contained 100 nM pre-activated SVMP, 200 nM human prothrombin and 0.5 mM S-2238 in reaction buffer. Assays were monitored as stated above, with reaction rates being determined in GraphPad (Prism) from the slope of the initial linear portion of each progress curve. Statistical significance was determined by one-way ANOVA followed by Dunnett’s multiple-comparisons test.

### Plasma clotting assay

The research involving derivatives of human blood samples was approved by the United Bristol Healthcare NHS Trust research ethics committee, project E5736, and all participants provided informed consent in line with the Declaration of Helsinki. Blood was collected from healthy adult volunteers not taking platelet-affecting medications and without bleeding disorders, pregnancy, or blood-borne infections. Samples were drawn into 3.2% sodium-citrate Vacutainers, and platelet-rich plasma (PRP) was prepared by centrifuging citrated whole blood at 180 × g for 17 minutes at room temperature.

Prior to the plasma clotting assay, SVMPs were activated as follows: 1 μM of each construct was incubated with 750 μM ZnCl in 50 mM HEPES pH 7, and 150 mM NaCl overnight, at 37°C.

To assess the SVMPs-induced coagulation, we performed an absorbance-based plasma clotting assay as described (Still *et al*, 2017), with some modifications. Briefly, 10 μL of sample (200, 100, 50 and 25 nM final concentration) was added to each well in a clear half-area microplate. Next, 20 μL of 50 mM CaCl_2_ was added, followed by 20 μL of PRP. The plate was then immediately read for kinetic absorbance (every 2 minutes) at 595 nm for 90 minutes using a TECAN Infinite M200 Pro plate reader.

Assays were performed twice, with each assay containing technical duplicates. The maximum clotting velocity of each of the curves was calculated as per clot waveform analysis (Sevenet & Depasse, 2017), by calculating the maximum of the first derivative, using GraphPad Prism. Multiple comparisons one-way ANOVA test was used to compare the maximum clotting velocity for each construct, using GraphPad Prism.

### Cytotoxicity assay

MTT assay was based on the methods of Hall *et al*. (*47*). HUVEC cells (HUVEC-c pooled, PromoCell) were seeded (5,000 cells/well, clear-sided microplates (SARSTEDT)) in complete medium (Endothelial Cell Growth Medium 2, PromoCell) and incubated for 24 hours at 37°C, 5% CO_2_. The cell line was freshly procured and checked for contamination before the start of the experiment. The next day, the SVMPs treatments (200, 400 and 800 nM) were prepared in complete medium, and cells were treated with each prepared solution (100 µl/well, duplicate wells) for 24 hours. For this assay, the MTT Cell Viability Assay Kit (Biotium) was used: 10 μL of MTT solution was added to the 100 μL of medium in each well and mixed by tapping gently on the side of the tray. The cells were incubated at 37°C for 4 hours. 200 μL DMSO was added directly into the medium in each well and pipetted up and down several times to dissolve the formazan salt. The absorbance was measured on BioTek Synergy Neo2 spectrophotometer at 570 nm. The % of cell viability for each treatment well was calculated as follows with buffer-treated cells as a control (100% viability):

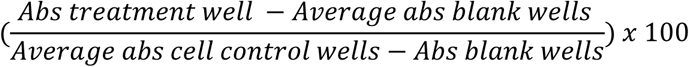

## Materials availability

Materials from this study are available from the corresponding authors upon reasonable request.

## Acknowledgements

The authors thank all members of the Berger, Schaffitzel and Casewell teams, past and present, for discussions and support.

## Additional Information

Competing interests: IB is shareholder and director of Geneva Biotech SARL commercialising the MultiBac system. The other authors declare that they have no competing interests

## Author contributions

Sophie Hall, Conceptualization, Data curation, Formal analysis, Investigation, Visualization, Methodology, Writing – original draft; Bronwyn Rand, Data curation, Formal analysis, Investigation, Visualization, Methodology, Writing – original draft; Iara Aime Cardoso, Data curation, Formal analysis, Methodology; Adam Robinson, Data curation, Investigation; Mark C Wilkinson, Data curation, Formal analysis, Investigation, Methodology; Dakang Shen, Sabastian Fernandes, Georgia Balchin, Konrad Kamil Hus - Methodology, Writing - review and editing; Nicholas R Casewell, Alastair Poole, Supervision, Writing – review and editing; Imre Berger, Conceptualization, Supervision, Methodology, Writing – original draft; Christiane Schaffitzel, Conceptualization, Supervision, Methodology, Writing – original draft, Writing – review and editing.

## Data availability

All data generated or analysed during this study are included in the manuscript and supporting files.

## Supplementary Figures

**Supplementary Figure 1:**
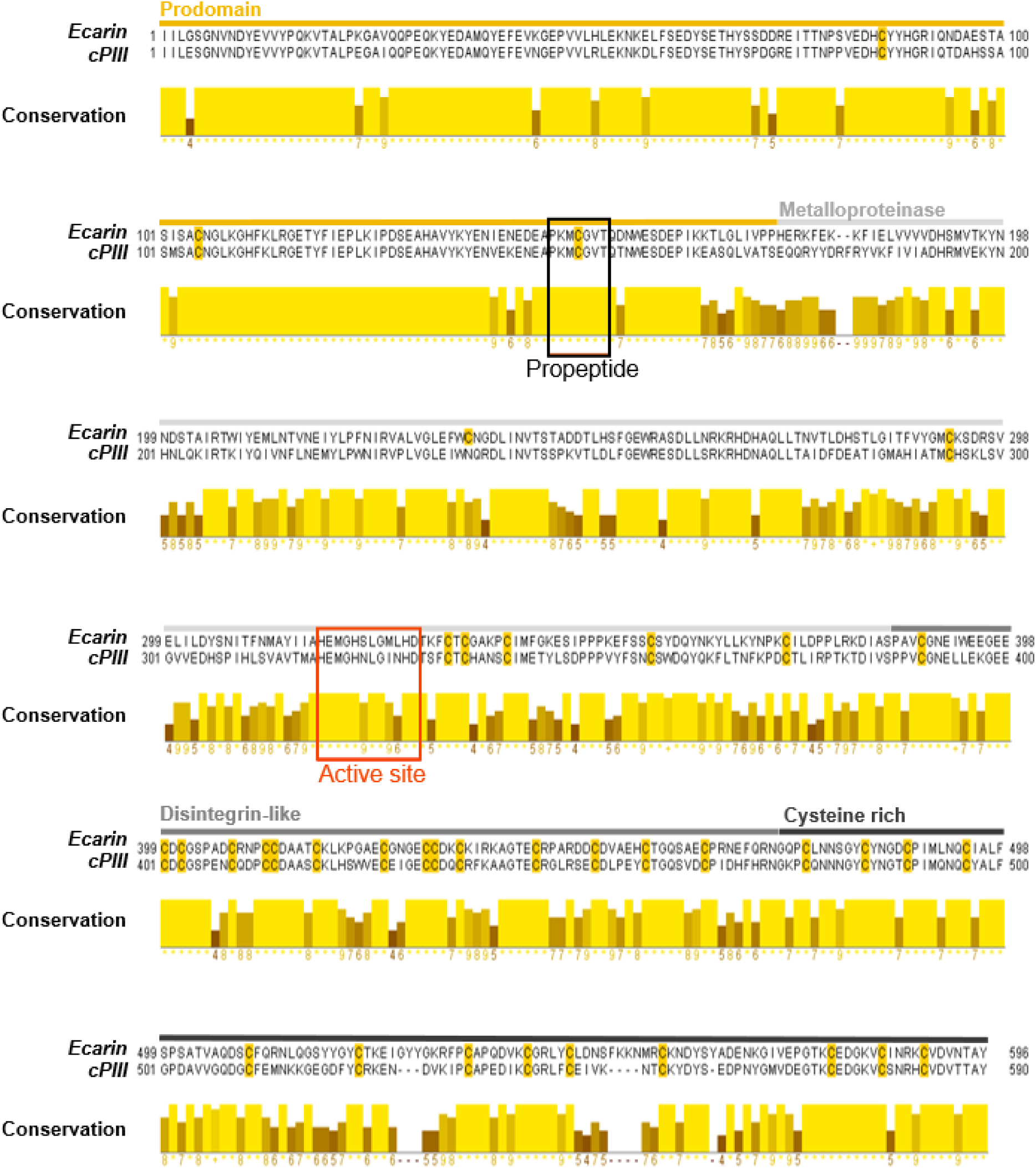
**Multiple sequence alignment of cPIII and Ecarin zymogen sequences**. Prodomain (orange), metalloproteinase domain (light grey), disintegrin-like domain (grey) and cysteine-rich domain (dark grey) domain boundaries are indicated. Black box: 7-amino acid propeptide; orange-red box: active site. Cysteine residues are highlighted in orange in the sequence alignment. Percentage identity: 65.05% (NCBI BLASTp (Camacho *et al*, 2009)).

**Supplementary Figure 2.**
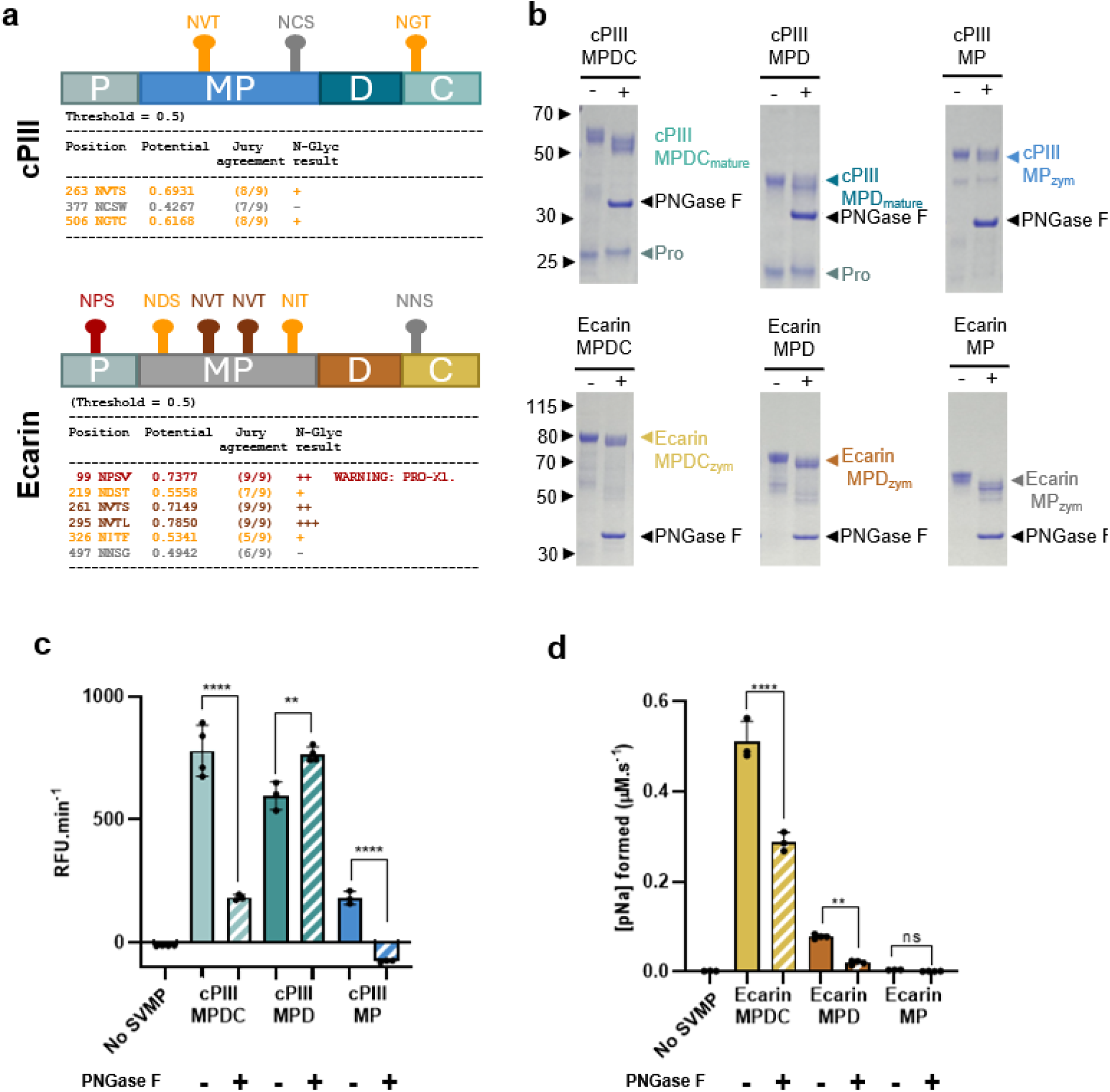
Impact of glycosylation on SVMP activity. (**a**) Predicted N-glycosylation sites in full-length cPIII (top) and Ecarin (bottom) identified using NetNGlyc 1.0. Domain orgnaisation is shown in a schematic view. MP stands for metalloproteinase, D for Dis and C for C-rich domains. Predicted glycosylation sites are indicated (colored according to probability scores shown in tables below). (**b**) Reducing SDS-PAGE analysis of purified SVMP zymogens before and after PNGase F treatment. Deglycosylation resulted in a shift to lower molecular weight for cPIII MPDC, cPIIlI MPD, and all Ecarin constructs. Activity of (**c**) cPIII constructs towards the fluorogenic substrate ES010 and (**d**) Ecarin constructs towards the chromogenic substrate S-2238, before and after treatment with PNGase F. Statistical significance was determined by one-way ANOVA followed by Šidák’s multiple-comparisons test.

**Supplementary Figure 3:**
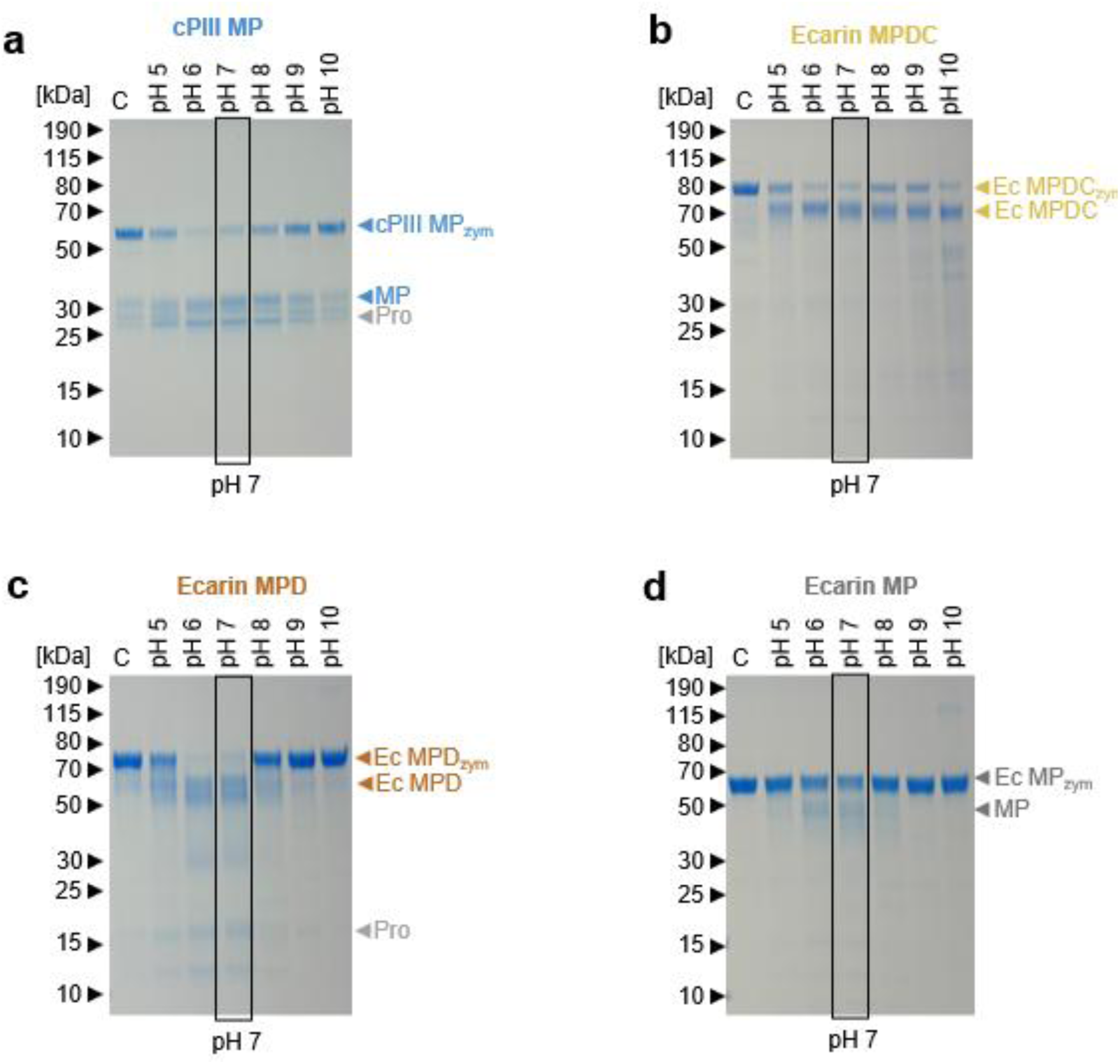
**pH screen for activation of recombinant zymogens to mature protein**. For activity assays, (**a**) cPIII MP, (**b**) Ecarin MPDC, (**c**) Ecarin MPD and (**d**) Ecarin MP zymogens were activated in buffer at pH 7.

**Supplementary Figure 4:**
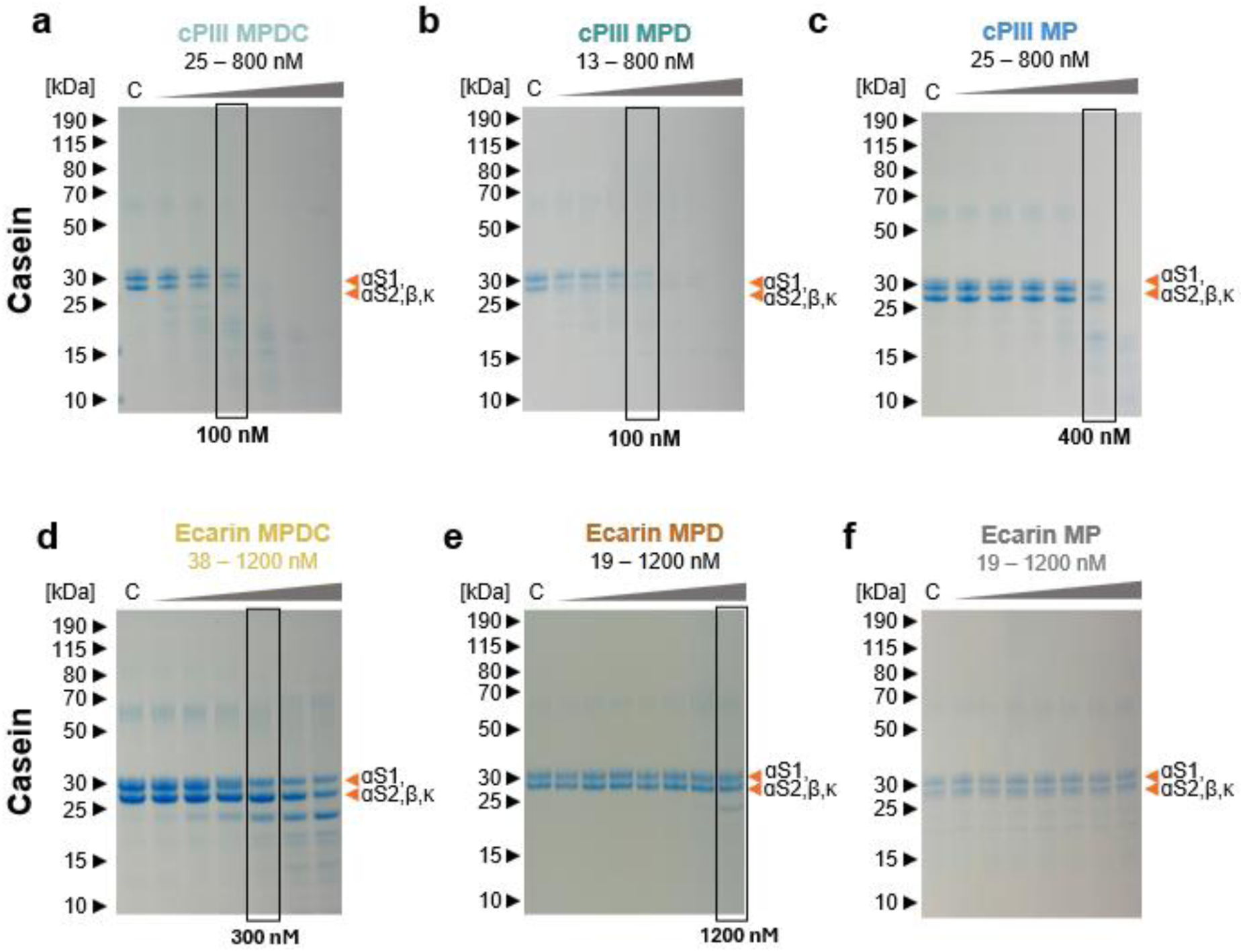
Non-specific *in vitro* SVMP activity assays. Casein degradation in the presence of a serial dilution of (**a**) cPIII MPDC, (**b**) cPIII MPD (**c**) cPIII MP, (**d**) Ecarin MPDC, (**e**) Ecarin MPD and (**f**) Ecarin MP SVMPs. Orange arrowheads: domains αS1, αS2, β and κ. C (control): pre-incubation of SVMP for 30 minutes in the presence of 20 mM EDTA.

**Supplementary Figure 5:**
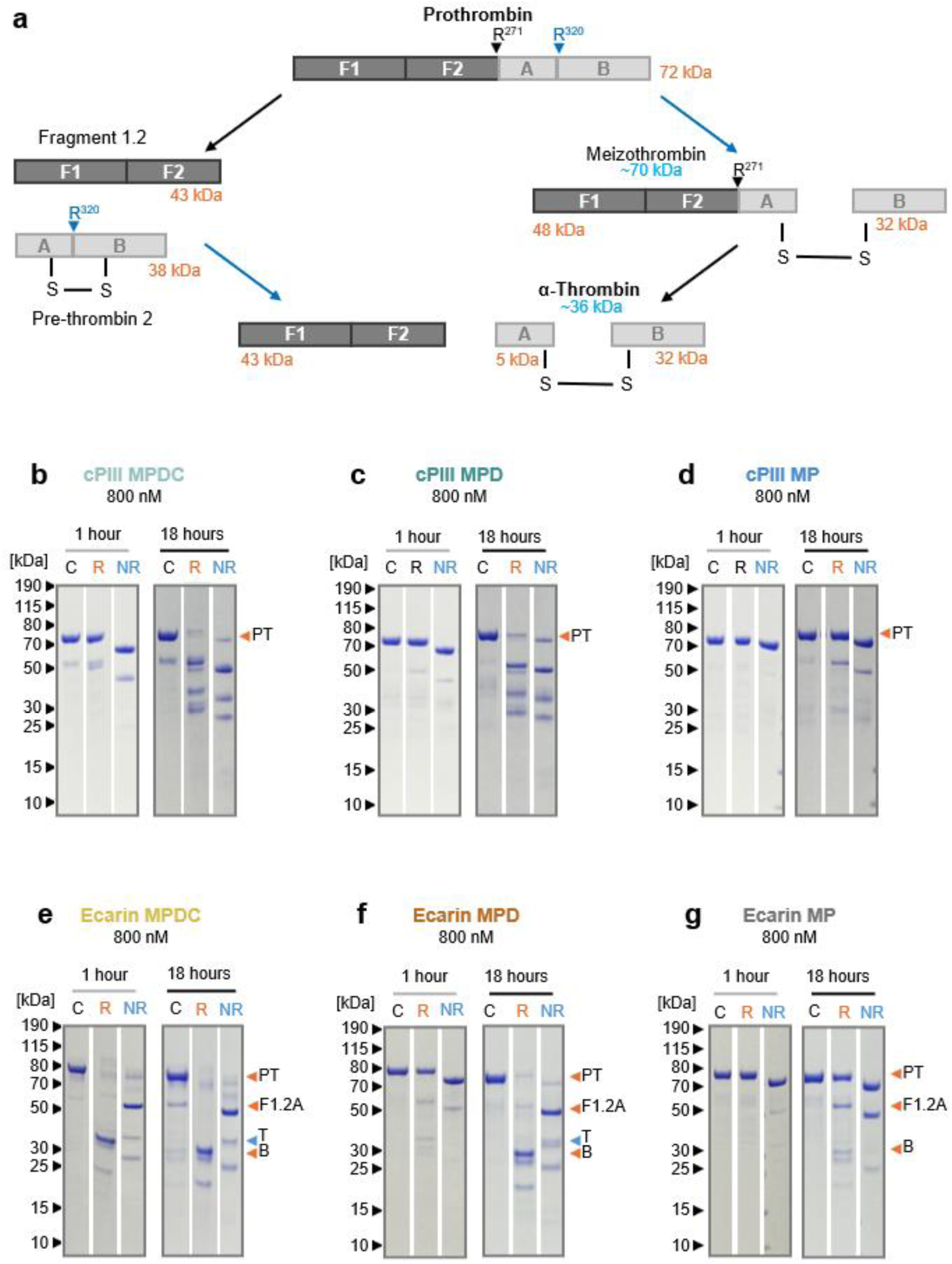
Reducing (R) and non-reducing (NR) SDS-PAGE analysis of prothrombin activation by SVMPs. (**a**) Schematic representations of PT activation, with molecular weight sizes as expected with reducing (orange) and non-reducing (blue) SDS-PAGE. PT activation in the presence of the serine-protease inhibitor, AEBSF, for 1 hour and 18 hours with 800 nM (**b**) cPIII MPDC, (**c**) cPIII MPD, (**d**) cPIII MP, (**e**) Ecarin MPDC, (**f**) Ecarin MPD and (**g**) Ecarin MP. The ∼30 kDa band (B chain of α-thrombin, B) shifts up to ∼36 kDa (A + B chains), confirming cleavage of prothrombin into thrombin (T) by Ecarin MPDC within (**e**)1 hour, and (**f**) Ecarin MPD in 18 hours. No thrombin band was observed for PT activation by the other SVMP constructs (**a**-**c, g**). C (control): prothrombin, with pre-incubated SVMP and 20 mM EDTA.

**Supplementary Figure 6.**
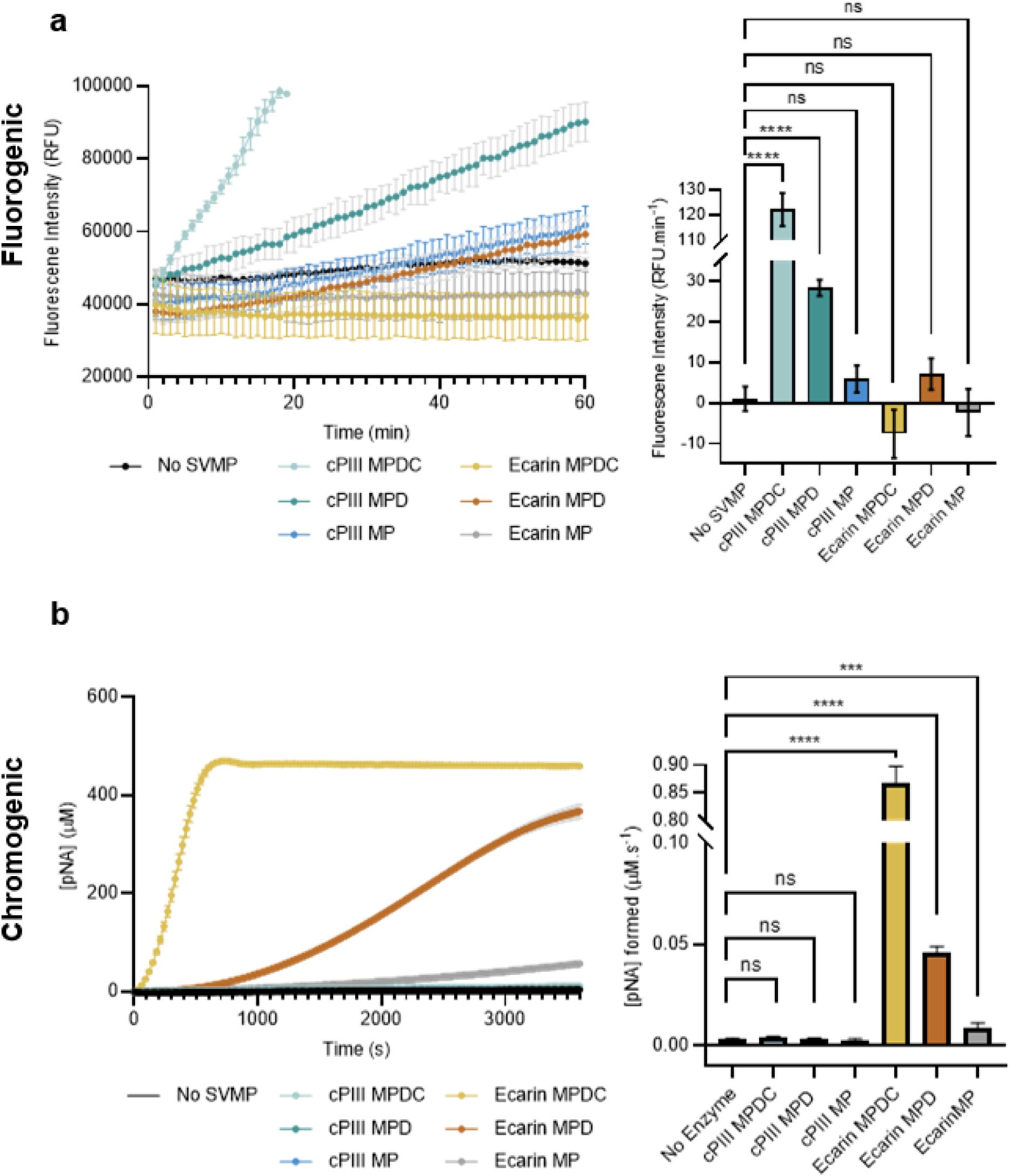
Fluorogenic and chromogenic assays of non-specific construct activity. (**a**) Raw fluorescence traces for ES010 cleavage by all constructs (left) and corresponding reaction rates calculated from linear regression of the initial linear phase (right). (**b**) Zeroed absorbance traces for S-2238 cleavage by all constructs (left) and corresponding reaction rates calculated from linear regression of the initial linear phase (right). Statistical significance was determined by one-way ANOVA followed by Dunnett’s multiple-comparisons test.

**Supplementary Figure 7.**
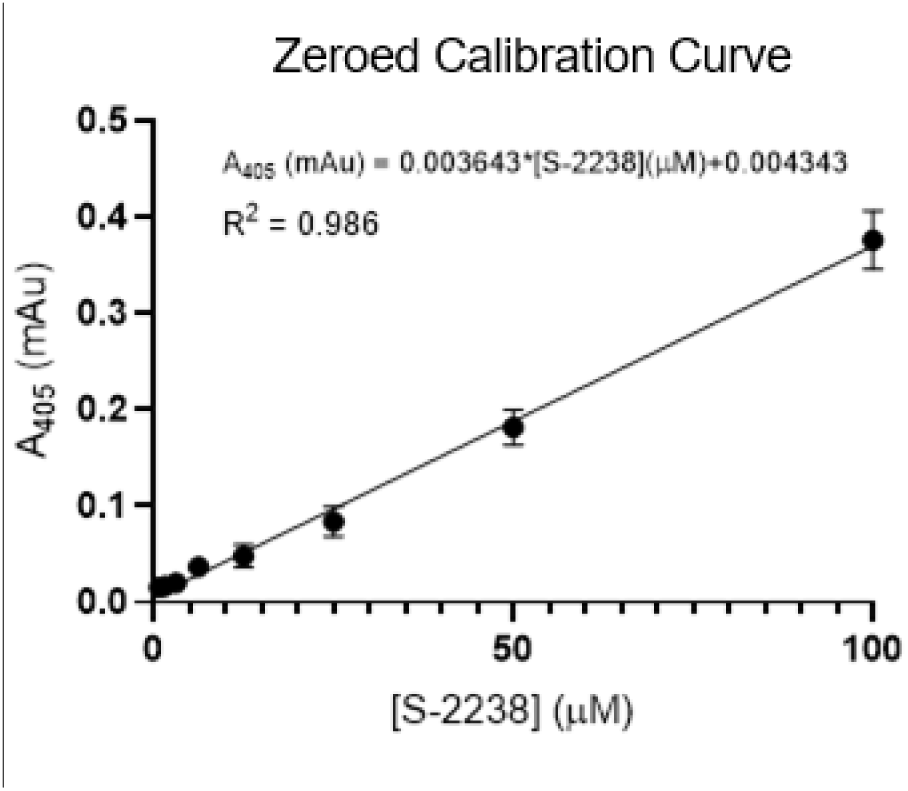
Calibration curve for S-2238 product quantification. Plateau absorbance values at 405 nm (A_405_) obtained following complete conversion of S-2238 to p-nitroaniline (pNA) were plotted against substrate concentration. The resulting linear regression was used to convert absorbance values to pNA concentration in S-2238 activity assays.

